# Computational redesign of TALE proteins for DNA-templated assembly of protein fibers

**DOI:** 10.1101/2024.10.21.619424

**Authors:** Robbert J. de Haas, Mark D. Langowski, Andrew J. Borst, Visakh V. S. Pillai, Gwendolyn E. Hoffmann, Martin Bongers, Suna Cheng, Catherine Treichel, Elizabeth M. Leaf, Mengyu Wu, Eric M. Lynch, Justin M. Kollman, Francesco S. Ruggeri, Carl Walkey, Renko de Vries, Neil P. King

## Abstract

Many viral proteins self-assemble into capsid structures, often using their genetic material as a template for assembly. To date, *de novo* designed capsid-like proteins do not require genetic material as a template for assembly, which can be both an advantage and a disadvantage depending on the use case. Templates are indispensable, for example, in the assembly of linear structures with well-defined lengths. As a first step towards fully *de novo* designed templated assembly, here we redesign proteins from the Transcription activator-like effector (TALE) family of transcriptional regulators to polymerize on double-stranded DNA (dsDNA) templates. Starting from natural TALE protein sequences, we created idealized repeat proteins with sequence-independent DNA binding properties that cooperatively self-assemble to form linear protein-DNA complexes with template-controlled lengths. We used high-resolution atomic force microscopy (AFM) and cryo electron microscopy (cryo-EM) to characterize the three-dimensional structures of the DNA-protein hybrid complexes. In these structures, a protein filament helically wraps around the dsDNA using a binding mode similar to that of natural TALE proteins. As an example application of these materials, we show the system can be used for repetitive peptide antigen display at precisely controlled repeat distances, and that such immunogens elicit robust antigen-specific antibodies in mice.

## Introduction

Viruses self-assemble into highly ordered nanoscale architectures with high fidelity, despite the presence of a complex background of host biomacromolecules and widely varying assembly conditions. One particularly effective strategy to ensure efficient encapsulation under these challenging and varying conditions is to use the virus genome as a blueprint to guide the assembly process^1^. For example, the Tobacco Mosaic Virus capsid proteins nucleate via a packaging signal in its RNA genome, which initiates a highly ordered RNA-templated assembly process^2,3^.

To date, the strategy of cargo-templated assembly has not yet been fully exploited in designing artificial capsid systems composed of *de novo* proteins^4^. While there are many use cases for artificial capsids that are not cargo-templated, one can also imagine many use cases where cargo-templated assembly would be beneficial. For example, protein-encapsulated dsDNA could be useful as a vaccine, with the protein serving as a multivalent antigen display scaffold^5^ and the cargo potentially acting as an adjuvant through activation of Toll-like receptors (TLRs)^6^. Another use case could be nucleic acid delivery^7,8^. Finally, from a nanomaterials design perspective, templated assembly is one of the few established mechanisms for achieving assembly into linear structures of well-defined lengths^9–12^.

Minimal DNA-binding coat proteins have been designed previously^10^, but these were based on simple polypeptide domains that are less engineerable. A number of different naturally occurring protein families have evolved to bind and in some cases assemble on dsDNA^12,13^, and some of these are attractive building blocks for testing novel design strategies. For example, the TALE proteins are ideal scaffolds for two reasons. First, they have a quasi-repetitive core consisting of well-defined repeat units that could serve as the basis for idealized repeat proteins. Equally important, the interfaces between the repeated motif within each subunit could be repurposed to act as an interface that drives self-assembly of multiple subunits to form a polymer. Second, TALE proteins are known to undergo a slight allosteric transition upon DNA-binding^14,15^, a mechanism that could be exploited to favor assembly on the DNA template and to prevent assembly in the absence of DNA templates.

Here we leveraged these features of TALE proteins to computationally design idealized repeat proteins with sequence-independent DNA binding properties. We found that the proteins self-assembled on dsDNA templates to form highly ordered linear polymers that can be used to display peptide motifs with precisely controlled spacing.

## Results

### Design of sequence-independent DNA-templated protein polymers

We began from the known sequences and structures of sequence-specific TALE DNA-binding proteins. TALEs comprise three simple structural domains: an N-terminal region (NTR), a central repeat domain (CRD), and a C-terminal region (CTR) (**Fig. 1a**). The NTR has been suggested to serve as the indispensable nucleation site for DNA binding^16^, while the CRD consists of a set of quasi-repetitive 33–35-residue motifs^17^. The CRD repeats follow the major groove of dsDNA and form sequence-specific interactions solely through two adjacent residues (called the repeat-variable diresidue, RVD) located in the turn of the helix hairpin. Diverse combinations of RVD confer a selective preference for binding to nucleotides A, C, T, or G^18^. Owing to their modular, repetitive structures and simple rules for DNA-binding specificity, we hypothesized CRD repeats would make an excellent starting point for the design of repetitive sequence-independent DNA-binding proteins. We selected TALE PthoX1 as a starting point for design, since a high-resolution crystal structure was available (**Fig. 1a**; PDB ID: 3UGM)^17^.

**Figure 1.**
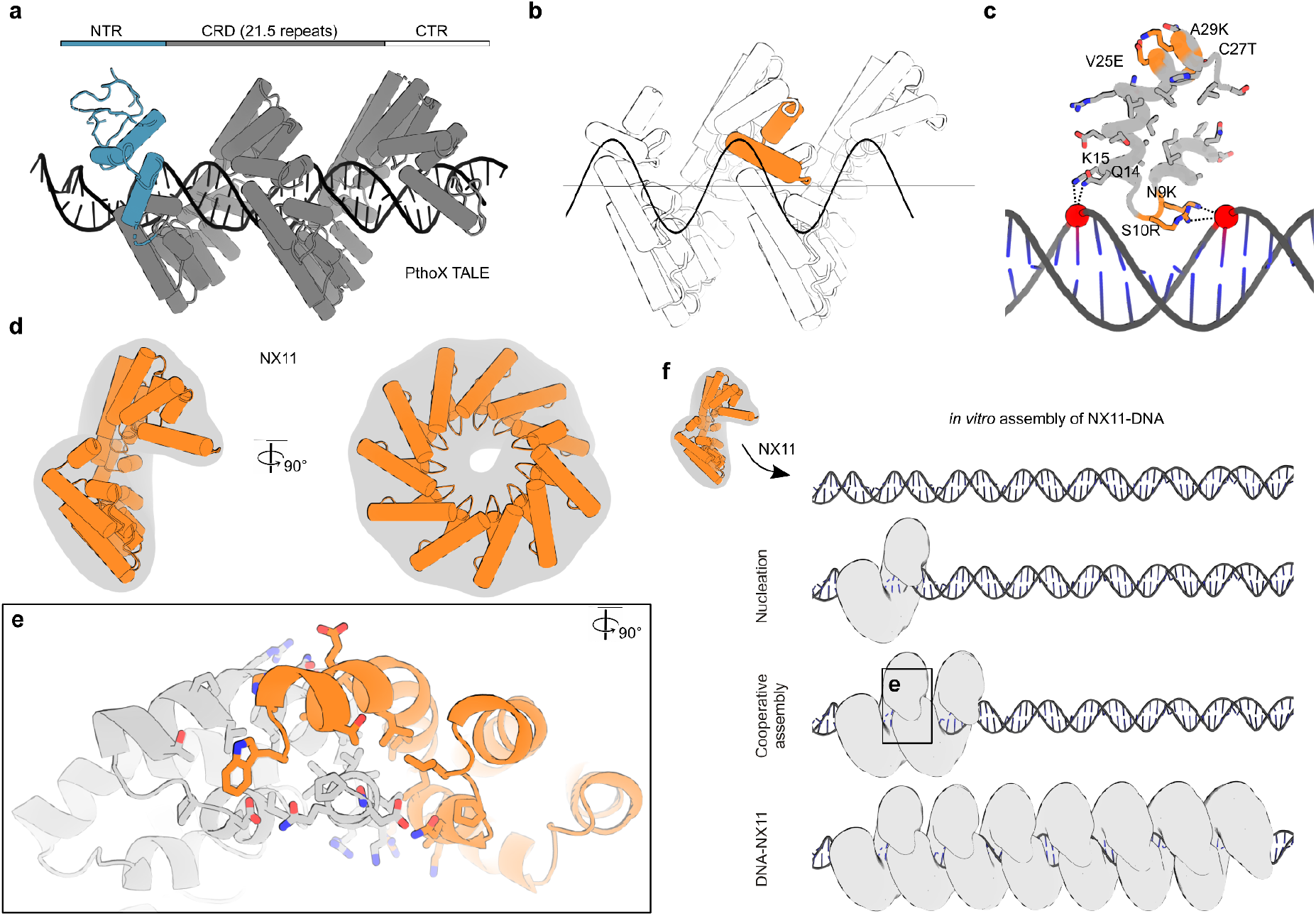
Design of sequence-independent DNA-templated protein polymers. **(a)** PthoX Transcription Activator-like Effector Protein (TALE). *Top:* domain architecture. NTR, N-Terminal Region; CRD, Central Repeat Domain containing helix-turn-helix repeats where each CRD repeat binds to a specific DNA base pair; CTR, C-Terminal Region. *Bottom:* Cylinder cartoon diagram of the crystal structure of PthoX (PDB ID: 3UGM^17^). Only a part of the NTR (blue) and the full CRD (gray) were crystallized. **(b)** A single CRD repeat is extracted, aligned to an idealized dsDNA strand, and helically symmetrized following the helical pitch of DNA of 34 Å and 10.5 monomers per turn. **(c)** Designed monomer, NX1, with symmetrical mutations indicated. S9R and N10K remove specificity and introduce additional non-specific interactions through ionic interactions with the phosphate backbone (red spheres) of DNA. Cys27Thr removes potential dimerization while Val25Glu and Ala29Lys mutate hydrophobic surface residues to a stabilizing salt-bridge. **(d)** Structure of NX11, propagated from NX1 repeats. **(e)** NX-NX intra-repeat interface formed by two adjacent NX proteins. This is the same interface as the inter-repeat interface and is mostly hydrophobic. **(f)** Hypothesized *in vitro* assembly of NX11 with DNA to form NX11-DNA complexes. Upon mixing, random nucleation occurs due to non-specific interactions between NX11 and DNA, followed by cooperative assembly starting from the nucleation site due to additional NX11-NX11 protein interfaces that form with multiple adjacent NX11 proteins (see **e**). Assembly continues until full coverage of the DNA is achieved, resulting in NX11-DNA complexes.

We extracted the 7th CRD 33-residue repeat (residues 498 to 527) because it already had a deletion of one of the RVD sites. We symmetrized it following the 35 Å superhelical pitch and 10.5 monomers per turn (one repeat per base pair) of idealized B-form DNA (**Fig. 1b**). We then used Rosetta to symmetrically re-design the monomer (see Methods). Residues 9 and 10 on the helix preceding the RVD site were re-designed with constrained backbone relaxation to facilitate minor backbone movements, while the native non-specific DNA-interacting residues Gln14 and Lys15 (ref. ^19^) were restricted from repacking. Rosetta suggested Ser9Lys and Asn10Arg mutations could contribute to non-specific DNA binding via ionic interactions with phosphate groups in the DNA backbone (**Fig. 1c**). We also conservatively redesigned hydrophobic residues on the monomer surface to potentially improve its stability and solubility. Cys27Thr was included to remove potentially problematic disulfide bond formation, and Val25Glu and Ala29Lys introduced a stabilizing salt bridge in place of surface hydrophobes (**Fig. 1c**).

Next, we used Rosetta Remodel^20^ to fuse position 34 of one symmetrically arranged monomer to position 1 of its neighboring monomer (see Methods). Repeat proteins with defined numbers of repeats were generated by symmetrically propagating the backbone of the monomer. We refer to the resultant sequence-independent DNA-binding repeat proteins as NucleoX, or NX, followed by a number referring to the number of repeats present (e.g., NX11 has 11 repeats) (**Fig. 1d**).

The designed NX proteins, like the naturally occurring TALE proteins, have predominantly hydrophobic interfaces between connected CRD repeats (**Fig. 1e**). Unlike the TALE proteins however, which have capping CTR and NTR domains, the NX proteins have exposed CRD repeat interfaces at their N and C termini. We hypothesized the CRD would allow for interactions between neighboring NX proteins that would drive self-assembly and cooperative DNA binding (**Fig. 1f**). This hypothesis assumed that the non-specific protein-DNA interactions we engineered into the NX proteins would be sufficient to nucleate their assembly on DNA, even in the absence of the NTR domain that has this function in the natural TALE proteins^15^. Furthermore, because TALE proteins are known to undergo a conformational change upon binding to DNA^21,22^, we assumed that the structure of the intermolecular interfaces would be slightly different in solution, which would disfavor assembly prior to DNA binding.

### Assembly with DNA and nuclease protection

To facilitate purification, we generated expression plasmids encoding NX5 and NX11 with a cleavable hexahistidine tag at the N terminus. Potential steric hindrance in the protein-protein interface was minimized by eliminating the two terminal loop residues of the final repeat (Thr and Leu). Finally, a sole Trp residue was affixed to enable protein quantification via UV/vis spectroscopy. We expressed the NX5 and NX11 proteins in *E. coli*. Both constructs expressed well and were purified using immobilized metal affinity chromatography (IMAC) (**Fig. S1**). For simplicity we will focus here on NX11, but further biochemical and assembly characterization of NX5 can be found in the Supplementary Materials (**Fig. S2**). Importantly, NX11 eluted as a monomer from size-exclusion chromatography (SEC) (**Fig. 2a**), showing no indication of oligomerization in the absence of a DNA template. Circular dichroism (CD) performed on SEC-purified protein indicated a strong α-helical signal that was lost at 95 °C and completely regained after cooling, suggesting that the protein adopted the target α-helical structure (**Fig. 2b**).

**Figure 2.**
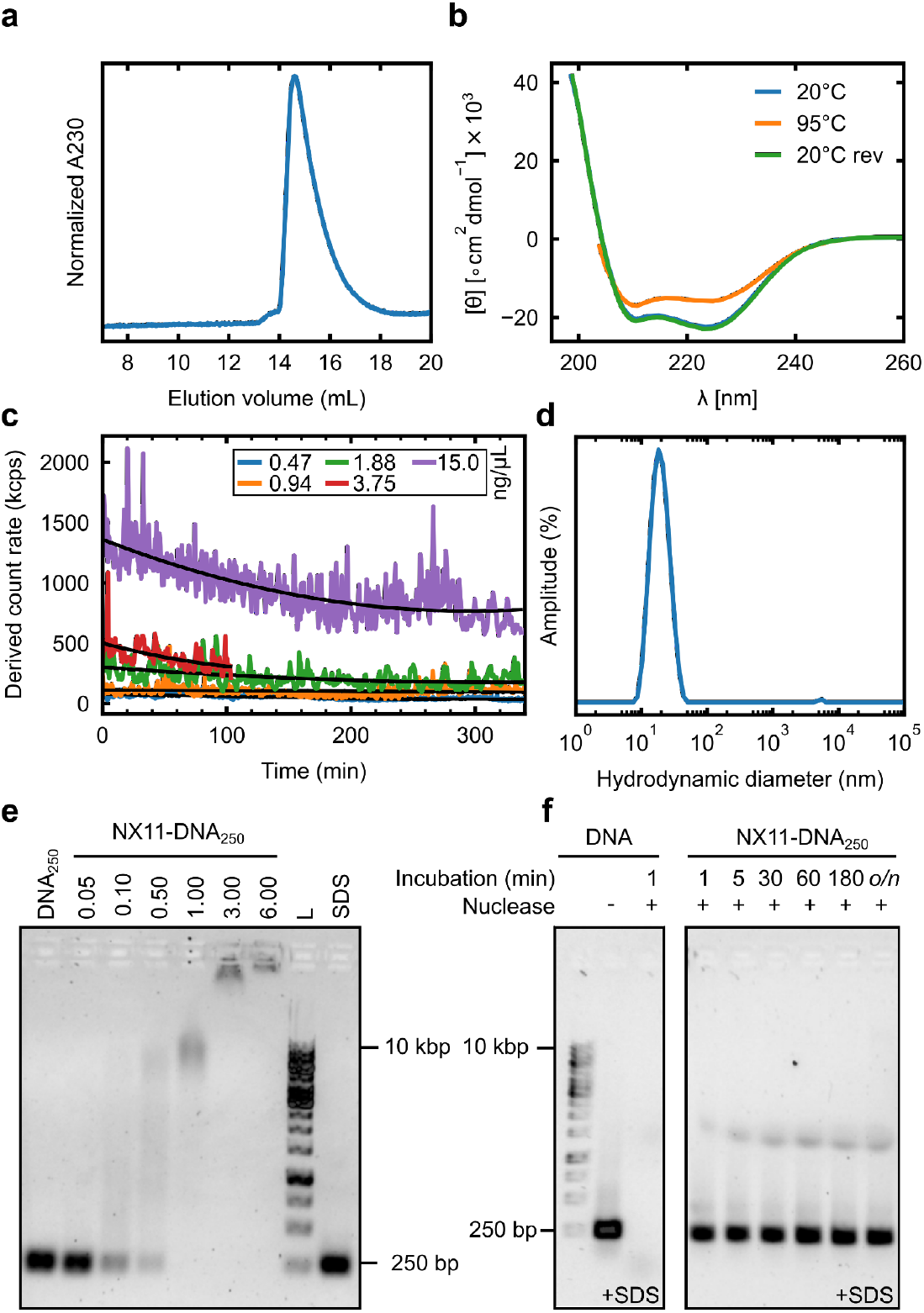
Assembly with DNA and nuclease protection. **(a)** Size-exclusion chromatography on Superdex 200 10/300 (Cytiva) demonstrating elution of monomeric NX11 protein at ∼15 mL, corresponding to a molecular weight of ∼42 kDa. **(b)** Circular dichroism of 0.1 mg/mL purified NX11 in PBS. Molar residue ellipticity [θ] is plotted as a function of wavelength, showing a predominantly α-helical profile. Heating to 95 °C led to a minor change in ellipticity, and subsequent cooling to back to 20 °C (20 °C rev) gave the same profile as before, demonstrating that NX11 proteins are thermostable. **(c)** Assembly kinetics of NX11 with different concentrations of 250 bp DNA (DNA_250_), tracked with static light scattering. NX11-protein was kept at a 3-fold excess relative to the number of DNA_250_ binding sites (assuming 1 NX per bp). Derived count rate in kilo counts per second (kcps) was recorded over time as a measure of assembly. Upon mixing NX11 and DNA the scattering intensity increases rapidly followed by a gradual decrease. **(d)** Dynamic light scattering intensity vs. size of assembled NX11-DNA_48_ complexes in PBS following > 4 hr incubation at room temperature. The average hydrodynamic diameter was 20 ± 6.5 nm. **(e)** Electrophoretic mobility shift assay for detecting assembly of NX11 with DNA_250_. NX11 was added at different ratios of excess. Due to charge neutralization and increase of the DNA-protein complex size, the DNA migrates less far into the gel, indicating complex formation. L is a 1 kb Gene ruler (Thermo Fisher Scientific), SDS is NX11-DNA complex loaded in presence of ∼0.2% SDS in the loading dye to denature the complexes. **(f)** Nuclease protection assay. The gel was run under denaturing conditions (∼ 0.2% SDS in loading dye) to clearly visualize the DNA. *Left:* control free DNA was fully degraded in < 1 min by Benzonase nuclease. *Right:* NX11-DNA complexes show full-length DNA_250_ at all incubation times, indicating excellent DNA protection from Benzonase.

We used static light scattering to follow the co-assembly kinetics of NX11 with various concentrations of 250 bp DNA (DNA_250_), maintaining a 3-fold molar excess of NX11 to DNA binding sites (assuming 1 NX repeat per bp). At the highest DNA_250_ concentration (15 ng/μL), assembly with NX11 led to a rapid increase in scattering intensity in the time between mixing and measurement followed by a gradual decrease over ∼4 hr (**Fig. 2c**). This suggests a two-phase assembly, with a rapid nucleation of multiple NX11 molecules on DNA followed by a slow re-arrangement. Dynamic light scattering (DLS) of assembled NX11-DNA_48_ showed these assemblies had a hydrodynamic diameter of 20 ± 7 nm and a polydispersity index of 0.13 (**Fig. 2d**).

To ascertain whether the assembly of NX11 with DNA is indeed cooperative, we assembled NX11-DNA complexes at increasing protein concentrations for a fixed concentration of DNA_250_. Electrophoretic mobility shift assays (EMSA) showed that upon increasing the excess of NX11 to DNA, mobility was reduced (**Fig. 2e**). At roughly 3-fold excess, the assemblies did not migrate into the gel due to essentially complete charge neutralization. The reduction in band intensity at higher NX11:DNA ratios is probably the result of steric hindrance or distortion of the DNA structure caused by the NX11 protein preventing the binding of Sybr Safe dye to the DNA: when analyzing denatured NX11-DNA assemblies using ∼0.2% sodium dodecyl sulfate (SDS) surfactant in the loading dye, we observed full-length DNA_250_ with similar intensity as the control in all samples. A similar EMSA experiment for NX5 with DNA assembly also showed complex formation (**Fig. S2**).

If NX11 assembles on DNA as intended, it would form a continuous protein coat that would be expected to protect the DNA template against nuclease activity. To test this, we assembled NX11-DNA complexes and exposed them to Benzonase in an optimal cleavage buffer for various incubation times before adding 50 mM EDTA to chelate Mg^2+^ and Ca^2+^ co-factors and terminate the reaction. EMSA performed in the presence of ∼0.2% SDS revealed that full-length DNA_250_ could be observed even after overnight (>18 h) incubation with Benzonase, while the control DNA was fully cleaved in less than 1 minute (**Fig. 2f**). These data indicate that NX11 indeed forms a tightly bound protective coating on the DNA template.

Next, to explore the importance of the NX11-NX11 interface for DNA coating and protection, we performed an alanine scan to identify key residues contributing to the interface strength (see Methods). We successfully purified a single-alanine mutant, NX11-1A (V25A), and a triple-alanine mutant, NX11-3A (V25A, L36A, L372A) (**Fig. S1**). EMSA of NX11-DNA_250_, NX11-1A-DNA_250_, and NX11-3A-DNA_250_ assemblies demonstrated that the alanine mutants form more intermediate assembly species, and that complete assembly required a larger excess of protein (**Fig. S3**). Benzonase challenge revealed that the mutant NX11-1A-DNA_250_ and NX11-3A-DNA_250_ complexes provided considerably less protection of their template compared to NX11 (**Fig. S4**). Together, these data show that an intact NX11-NX11 interface is required to drive complete and cooperative assembly of NX11-DNA complexes.

### Structure of NX11-DNA complexes

We used high-resolution Atomic Force Microscopy^23^ (AFM) to characterize the structure of NX11-DNA_750_ complexes as well as the DNA_750_ template alone (**Fig. 3a and Fig 3b)**. The average cross-sectional height of DNA_750_ and NX11-DNA_750_ complexes obtained from AFM were 0.50 ± 0.05 nm and 1.85 ± 0.40 nm, respectively (**Fig. 3c**). These values are consistent with the expected values for cross-sectional diameters of ∼2 nm for DNA and ∼3 nm for NX11-DNA complexes due to the non-linear underestimation that occurs in AFM when measuring cylindrical objects with diameters smaller than ∼4 nm^24^. Our quantitative single-molecule analysis demonstrated that the heights of NX11-DNA assemblies are significantly different from those of DNA_750_ alone (p value < 0.001), indicating that NX11 proteins are bound to the DNA. Analyzing contour lengths revealed an average contour length of 250 ± 20 nm for DNA_750_ (3.4 Å per bp) (**Fig. 3d**). Contour lengths of the NX11-DNA complexes were slightly shorter than those of DNA (240 ± 20 nm; p < 0.05). This result indicates some level of structural rearrangement of the DNA upon NX11 binding, resulting in condensation of the DNA along the long axis of the complexes.

**Figure 3.**
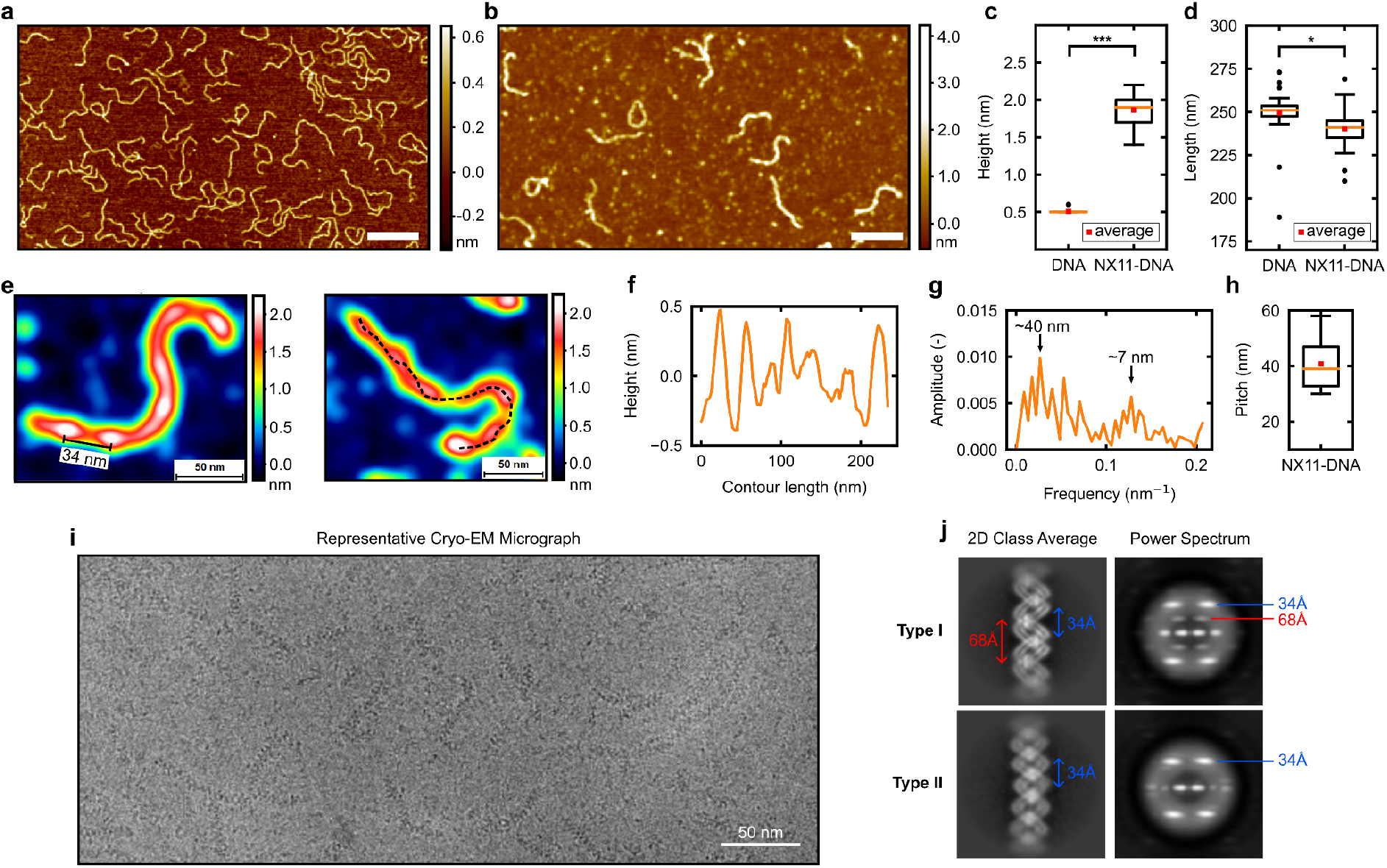
Structure of NX11-DNA complexes. **(a)** AFM map of DNA_750_. Scale bar: 200 nm. **(b)** AFM map of NX11-DNA_750_ complexes. Excess NX11 proteins are visible in the background. Scale bar: 200 nm. **(c)** Average cross-sectional height of individual DNA molecules (0.50 ± 0.05 nm) and NX11-DNA_750_ complexes (1.85 ± 0.40 nm). *** represents p-value < 0.001. The box represents the interquartile range (IQR) and the whiskers are Q1-1.5IQR and Q3+1.5IQR. **(d)** Average contour length of DNA (250 ± 20 nm) and NX11-DNA_750_ complexes (240 ± 20 nm). 35 particles were analyzed. * represents p-value < 0.05. The box represents the interquartile range (IQR) and the whiskers are Q1-1.5IQR and Q3+1.5IQR. **(e)** Zoomed in AFM images of representative individual NX11-DNA_750_ particles viewed by AFM. An example of a ∼40 nm observed periodicity is indicated in the figure. Dotted line represents cross-sectional profiles. **(f)** Representative cross-sectional height profile of a single NX11-DNA particle, measured by AFM. **(g)** Representative example of the Fast Fourier Transform of the first order derivative of a cross-sectional height profile. Two main peaks, corresponding to height periodicities on the NX11-DNA_750_ contours were observed at ∼40 nm, and ∼7 nm. **(h)** Analysis of 10 NX11-DNA_750_ complexes showing the distribution of the main periodicity (average: 40 ± 10 nm). The box represents the interquartile range (IQR) and the whiskers are Q1-1.5IQR and Q3+1.5IQR. **(i)** Representative cryo-EM micrograph of NX11^Q21.5^-DNA_48_ complexes. **(j)** Representative 2D class averages of two main types of complexes found (type I and type II). Type I appears to show a double-coiled assembly with apparent helical pitches of 34 Å and 68 Å, while type II only has the apparent 34 Å pitch. Pitches are determined from the X-shaped power spectra for each class average.

AFM analysis revealed that the NX11-DNA assemblies exhibited periodic fluctuations in height along the long axis (**Fig 3e**). We analyzed this periodicity at the single molecule level by acquiring height profiles along the main axis of symmetry of the NX11-DNA complexes. A first analysis of the cross-sectional profiles revealed a periodicity of ∼40 nm (**Fig. 3f**). An accurate and unbiased analysis of the periodicity of the profiles was obtained via Fast Fourier Transform (FFT), which confirmed the presence of a major periodicity of 40 ± 10 nm and additionally revealed a smaller periodicity of 7 ± 2 nm (**Fig. 3g,h and Fig. S5**). We speculate that the former is an effect of slight mismatch between the NX11 helical twist and the helical twist of the DNA, which potentially builds up frustration in the DNA double helix, ultimately leading to less well-defined areas of NX11 binding.

Negative stain electron microscopy (nsEM) was used to study smaller NX11-DNA complexes (<40 nm) in greater detail. When using short DNA templates (50 bp), we observed side-by-side dimerization of protein-DNA fibrils in 2D class averages. Such dimers were not observed for complexes in which longer templates (100 bp) were used (**Fig. S6**). Dimerization may be due to weak yet repetitive interactions along the DNA fiber. In an attempt to block such interactions, we introduced a Arg21Gln mutation on every second repeat of NX11, (NX11^21Q.5^). Indeed, no dimerization was detected during nsEM of NX11^21Q.5^-DNA_50_ fibers. The NX11^21Q.5^-DNA_50_ 2D class averages showed a continuous polymerization of protein on DNA as expected based on known structures of TALE-DNA complexes^17^ (**Fig. S7**).

To increase resolution, we analyzed NX11^21Q.5^-DNA_48_ complexes by cryo-EM (**Fig. 3i**). 2D class averages clearly demonstrated heterogeneity in the particle population. Two main types of NX11^21Q.5^-DNA_48_ complexes could be distinguished: Type I complexes appeared to show a double coil (**Fig. 3j**), with an additional double pitch of ∼68 Å relative to the expected ∼34 Å pitch in the design model. Type II complexes showed only the expected ∼34 Å in the class averages and a power spectrum analysis. Owing to heterogeneity within both complex types deriving from flexibility along the fiber axis, it was not possible to obtain accurate 3D reconstructions at high resolution. Nevertheless, AlphaFold2 predictions suggested that NX11^21Q.5^ could adopt an extended conformation with a helical pitch approaching 68 Å (**Fig. S8**), suggesting that the double-coil assembly may be formed by similar extended polymers.

In the absence of more detailed data, we conclude from the 2D class averages that NX11 proteins are closely packed and ordered on the DNA, suggesting that the designed protein-protein interfaces are formed. Additionally, no incomplete assemblies or assemblies of only NX11 were observed by either AFM or cryo-EM. Crucially, at high protein concentrations we demonstrated that NX11 does not oligomerize in solution by itself (**Fig. 2c**), suggesting that it is indeed a conformational change upon DNA binding that triggers the cooperative assembly of NX11.

### Antigen display and immunogenicity of NX11-DNA complexes

Ordered, DNA-templated NX11-DNA fibers could be used in a range of applications where having a nucleic acid cargo and a linear, repetitive structure could be advantageous. One such example is peptide antigen display for vaccine applications. The linear NX11-DNA structures should allow for presentation of peptide epitopes at high yet controlled density, with antigen copy number determined exactly by DNA template length.

To explore this application, we chose three peptide epitopes derived from the immunodominant repeat region of the *Plasmodium falciparum* circumsporozoite protein (PfCSP)^25^. Rosetta Remodel was used to model three and five copies of the major, minor, and junctional (junc) repeats of PfCSP inserted into the 3rd, 6th, and 10th surface loops of NX11 (majorX3, minorX5, etc.; **Table S2**). This arrangement would display the epitopes with roughly equal spacing on the DNA (**Fig. 4a**). All NX11 immunogens were successfully purified by IMAC and SEC. The proteins were assembled with highly pure pharmaceutical grade DNA_250_ (NoLimits, Thermo Fisher Scientific) into complexes at a 1.1× excess, and assembly was confirmed by EMSA (**Fig. S9**). nsEM micrographs showed NX11-DNA_250_ complexes were formed and appeared reasonably homogenous (**Fig. 4b**). DLS measurements of the complexes yielded hydrodynamic diameters of 20–30 nm, which is in line with rod-shaped particles of ∼6 nm diameter and ∼85 nm length (**Fig. 4c**).

**Figure 4.**
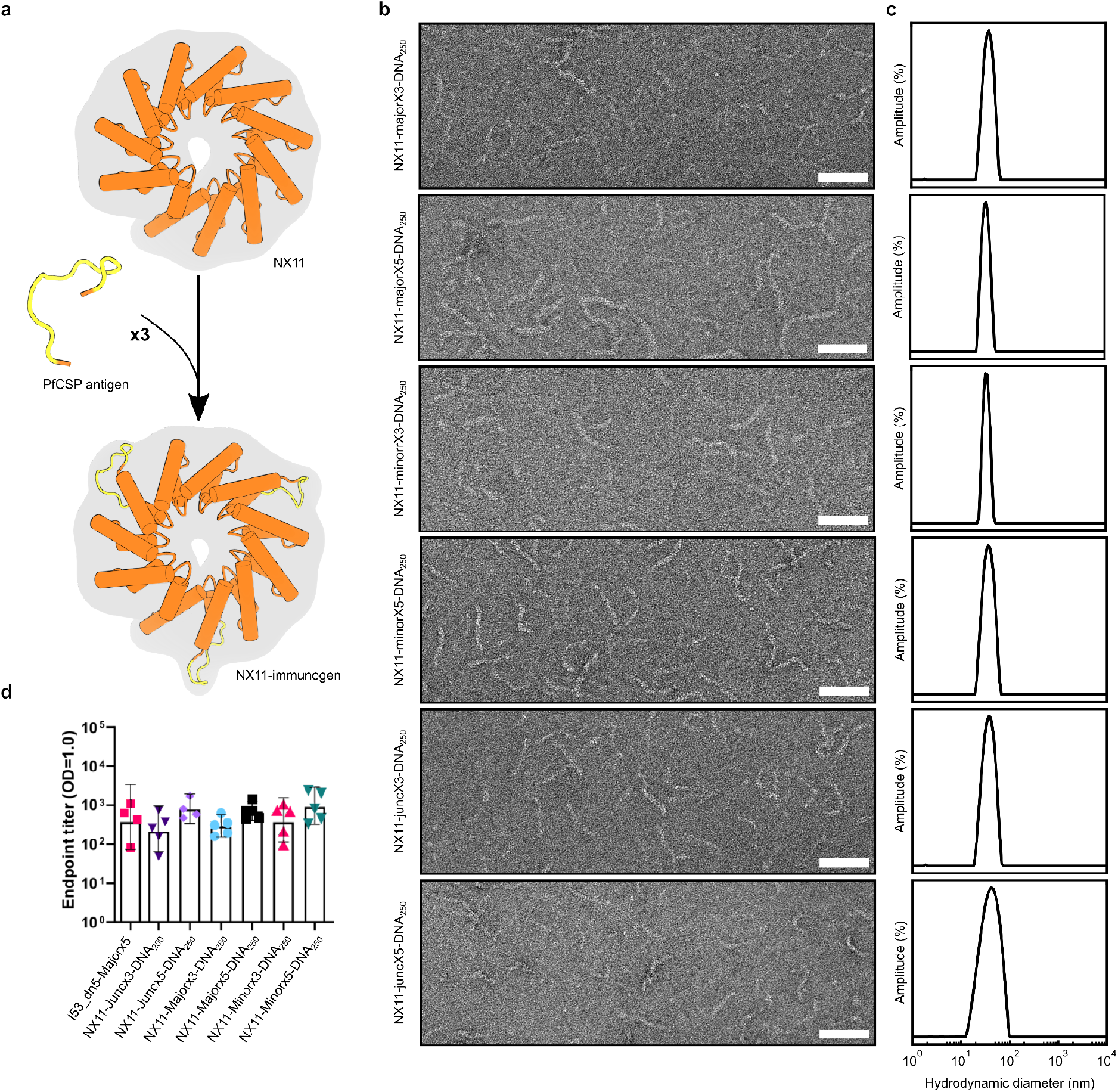
Antigen display and immunogenicity of NX11-DNA complexes. **(a)** Schematic of display of PfCSP tetrapeptide repeat epitopes at three locations in the NX11 protein. Repeats (x3 or x5) of major, minor, and junctional (“Junc”) epitopes were inserted in the surface-exposed loops of NX11. **(b)** Representative nsEM micrographs (scale bar = 100 nm) and **(c)** DLS of NX11-DNA_250_ immunogen complexes, showing a hydrodynamic diameter of ∼20-30 nm. **(d)** Endpoints titers of anti-PfCSP antibody in the sera of immunized mice (n=5 per group), measured by ELISA.

In previous work, computationally designed protein nanoparticles have been successfully used as multivalent display platforms for vaccine development^26–30^. To benchmark the performance of our linear protein-DNA immunogens against one of these, we engineered majorX5 into surface-exposed loops of the homopentameric component of a *de novo* two-component icosahedral nanoparticle (I53_dn5_majorX5), such that each particle displayed 60 copies of majorX5 antigen (**Fig. S10**).

We immunized groups of five B6(Cg)-*Tyrc-2J*/J mice intramuscularly with 1.5 µg of the NX11-DNA_250_ immunogens or the I53_dn5_majorX5 benchmark immunogen formulated in AddaVax at weeks 0 and 3. Serum collected two weeks post-boost was analyzed for PfCSP binding by enzyme-linked immunosorbent assay (ELISA), which showed antigen-specific antibody was elicited by all immunogens (**Fig. 4c**). Importantly, the NX11-DNA immunogens elicited similar antibody titers to the *de novo* icosahedral protein nanoparticle, indicating that the linear NX11-DNA platform shows potential as a multivalent vaccine platform (**Fig. 4d**).

## Discussion

In this study, we successfully redesigned a naturally occurring TALE protein to assemble into linear polymers on dsDNA templates. We leveraged the repetitive nature of the CRD domain by eliminating the naturally occurring capping domains and reusing the inter-repeat interface as a protein-protein interface. This approach is relatively straightforward and could be used to generate assemblies from other types of repeat proteins, which have proven to be versatile building blocks for designing new protein-based materials with high precision^31–34^. For instance, a similar strategy was recently employed to create protein building blocks that form a wide variety of architectures without extensive sampling of irregular interactions^31^. Our work extends this principle to templated assembly on nucleic acids, thereby providing a route to combining the accurate design of self-assembling proteins with DNA nanotechnology^35^.

Electron microscopy of NX11-DNA fibrils revealed a periodicity of ∼68 Å in addition to the expected pitch of ∼34 Å, which was confirmed here by AFM single-molecule analysis. Thus, EM and AFM suggested that two NX11 fibrils may wrap the DNA. The periodicity of around 65 Å is consistent with the AF2-predicted pitch for the unbound case, and there is experimental evidence suggesting that non-bound or non-specifically bound TALEs can adopt configurations with a longer superhelical pitch. For example, for the dHax3 TALE, the superhelical pitch increases from 35 Å in the DNA-bound form (PDB ID: 3V67) to ∼60 Å in DNA-free form (PDB ID: 3V6P)^22^. There is some evidence to suggest that TALEs first search DNA in a weakly bound long-pitch configuration, followed by a shortening of the pitch when binding more strongly to their target sequence^15,36^. We hypothesize that our NX11 proteins possess conformational flexibility similar to that observed in natural TALE proteins, which complicates high-resolution structural studies when bound to DNA.

As we demonstrated, linear protein-DNA complexes can be an interesting modality for the precisely controlled symmetrical display of antigens at high copy numbers. In earlier studies, designed self-assembling protein nanoparticles symmetrically displaying heterologous antigens proved highly immunogenic^28,29^, and we found that NX11-DNA–based immunogens were similarly immunogenic. In contrast to those finite (i.e., bounded) assemblies, NX11-DNA complexes of many different sizes and antigen copy numbers could be generated by using different protein variants or simply altering dsDNA template length. Linear display platforms characterized to date include simple peptide-based systems^7,10,37,38^ and complex assemblies based on filamentous viruses^39–41^. Due to their modular nature and the high density of exterior-facing loops, the NX11-DNA assemblies designed here provide precise and tunable control over antigen spacing, as well as the ability to include various DNA templates that could provide a beneficial adjuvanting effect^42^.

Lastly, we propose that variants of NX11 could be designed to ensure that nucleation occurs at specific DNA sequences, followed by elongation through the addition of sequence-independent binders or subunits. This refinement would enable more precise control over the assembly process and further functional optimization of the designed assemblies. In concept, this strategy could allow for the creation of programmable and addressable hierarchical assembly^43,44^ and complex asymmetrical protein assemblies that use DNA as a template^45^.

## Materials and Methods

### Computational design of NX-proteins

The 34 amino acid CRD protamer with the sequence PDQVVAIASNGGKQALETVQRLLPVLCQAHGLT was extracted from the PthoX1 crystal structure (PDB ID: 3UGM). The protamer was aligned to idealized B-form dsDNA^46^. Next, we enforced radial symmetry following the major groove of dsDNA (34 Å pitch, and 10.5 monomers per turn) and generated a symmetry definition file. Next, we used Rosetta XML-scripting for redesign. Existing native non-specific DNA interactions Gln14 and Lys15 were prevented from repacking. But other residues except for Gly and Pro near the RVD (and thus the DNA) were allowed to be symmetrically redesigned, with constrained backbone movements, resulting in S9K and N10R mutations. A second step of Rosetta design focused on optimizing idiosyncrasies in the monomer. Conservative Rosetta design of surface residues led to Cys27Thr, Val25Glu and Ala29Lys mutations. All other residues were repacked during design (except for DNA-interacting residues). The symmetrically arranged monomers were oligomerized by fusing loops at 34 and position 1 of the next monomer using Rosetta remodel^20^. Oligomers with repeat numbers of 5, and 11 repeats were generated by propagation of torsion angles. Final protein sequences are provided in **Table S1**.

For the design of NX11, displaying *pf*CSP immunogens (**Table S2**), the immunogens replaced surface-exposed loops (HGLT) of the NX repeats at positions at repeat numbers 3, 6 and 10 of NX11. Using Rosetta Remodel, immunogen consequences were inserted at 114-118, 231-235 and 345-349 of NX11, and two adjacent residues were also mutated to Gly to provide additional flexibility. Immunogen sequences were inserted and viability was tested by simulating 100 closures with Rosetta Remodel, while keeping the rest of the protein structure fixed. Final protein sequences are provided in **Table S1**.

### Protein expression and purification

Synthetic genes for individual components, each with an N-terminal TEV protease cleavage site hexahistidine purification tag (MGHHHHHHGSSENLYFQGS) and a C-terminal Trp to facilitate UV/vis quantification. Genes were codon optimized for *E. coli* expression and purchased as genes from Genscript ligated into the pET-29b(+) vector at the NdeI and XhoI restriction sites or as g-blocks from Twist Bioscience Corp. and assembled using Golden Gate Assembly Mix (New England Biolabs) into a modified pET-24(+) vector. The proteins were expressed in BL21(DE3) (New England Biolabs) in Luria Broth (10 g Tryptone, 10 g NaCl, 5 g yeast extract) in 2 L baffled shake flasks. Cells were grown at 37 °C to an OD_600_ ∼ 0.6, and induced with 1 mM Isopropyl ß-D-1-thiogalactopyranoside (IPTG; Sigma Aldrich). Expression temperature was reduced to 18 °C and the cells were shaken for ∼18 h. The cells were harvested and lysed by sonication using a Qsonica Q125 for 15 min with 2 s pulses at 80% amplitude in 50 mM Tris pH 8.0, 400 mM NaCl, 30 mM imidazole, 1% Triton X-100, 10% glycerol and 1 mM PMSF). Lysates were clarified by centrifugation at 30,000 *g* for 30 min and applied to a 5 mL column bed of Ni-NTA resin (Qiagen) for purification by IMAC. Resin was washed with 25 mL wash buffer I (50 mM Tris pH 8.0, 400 mM NaCl, 30 mM imidazole, 10% glycerol) or wash buffer I + 0.75% CHAPS for proteins used in animal studies to reduce endotoxins, followed by 25 mL wash buffer II (50 mM Tris pH 8.0, 2 M NaCl, 30 mM imidazole, 10% glycerol) and 25 mL wash buffer I. The protein of interest was eluted using a gradient of 50 mM Tris pH 8.0, 400 mM NaCl, 300 mM imidazole, 10% glycerol. During elution absorbance at 280 nm was monitored and ∼ 20 mL eluates were pooled and dialyzed against 5 L phosphate saline buffer (PBS) pH 7.4. Dialysate was concentrated in 10K MWCO centrifugal filters (Amicon) to 1.5 - 2 mg/mL, sterile filtered (0.22 μm) and applied to either a Superdex 200 Increase 10/300 (Cytiva), or Superdex 75 10/300 (GE Healthcare) using PBS pH 7.4 or TRIS-buffer saline (TBS) pH 8.0 as the running buffer. NX-immunogens used in mice study (i.e. NX11-majorX3 etc.) were tested for endotoxins levels < 2 EU/mg using a Limulus amebocyte lysate assay (Charles River).

### Circular Dichroism

Proteins were diluted to 0.15 mg/mL in PBS in a quartz cuvette (Hellma) with a 1 mm pathlength. On a JASCO J-715 (JASCO Corporation), spectral scans were averaged over 20 measurements with scan rate of 50 nm/min and a response time of 2 s. Scans were performed at 20 °C, 95 °C and after cooling back to 20 °C (20 °C rev). After reaching temperature the samples were incubated for > 10 min to equilibrate. Data with high tension above 600 V are not shown.

### Assembly of NX-DNA complexes

Pharmaceutical grade DNA (NoLimits, Thermo Fisher Scientific) — unless stated otherwise — assembled at 15 ng/µL with NX5 or NX11 proteins at 3x excess relative to binding sites on DNA (assuming 1 NX repeat per bp) in PBS pH 7.4 or TBS pH 8.0. The assembly was allowed to complete for > 4 hr at room temperature, and typically overnight at room temperature.

### Light scattering

NX11-DNA_250_ complexes were assembled in a 12 µl DLS quartz cuvette, and light scattering was recorded immediately. Light scattering was measured using a ZS-Nanosizer instrument (Malvern, UK) at a fixed angle of 173° and 20 °C. Measurements were averaged over 10 sec and the derived count rate (the raw scattering value corrected for X) was recorded using Zetasizer software version 7.13 (Malvern, UK). The ZS-Nanosizer instrument was also used to quantify the hydrodynamic diameter of NX11-DNA_48_ complexes using dynamic light scattering (DLS). Light scattering was measured at a fixed angle of 173 ° in a quartz cuvette at 20 °C. Hydrodynamic diameters were calculated based on the average of 50 measurements, with automatically determined optimal measuring settings by the Zetasizer software (version 7.13).

### Electron-mobility shift assay

A 1 % agarose gel containing 1X Sybr Safe staining (New England Biolabs) was cast. 15 µl NX11-DNA_250_ complexes were mixed with 3 µl 6X loading dye (Thermo Fisher Scientific). In the case of loading complexes under denaturing conditions, complexes mixed with 3 µl loading dye supplemented with 0.6% SDS (0.2% SDS final concentration). Agarose gel was run in 1X Tris-acetate-EDTA (Sigma Aldrich) buffer at 110 V for ∼30 min and imaged using a GelDoc imager (Bio-Rad).

### Nuclease protection assay

NX11-DNA_250_ assemblies (2.25 µg DNA_250_ in 150 µL) were formulated in a 1X reaction buffer (10 mM Tris pH 7.5, 2.5 mM MgCl_2_, 0.1 mM CaCl_2_), and 12.5 U Benzonase (Millipore, Cat. No. 70746) was incubated 1 min, 5 min, 30 min, 1 hr, 3 hr and overnight (∼18 hr). The reaction was stopped by adding 50 mM EDTA pH 7.4 to chelate Mg^2+^ and Ca^2+^. Complexes were mixed with loading dye (Thermo Fisher Scientific) supplemented with ∼0.2% SDS. Denatured complexes were run on a 1% agarose gel containing 1X Sybr Safe stain (New England Biolabs).

### Atomic Force Microscopy

DNA_750_ and NX11-DNA_750_ samples were prepared on positive functionalized mica substrates. To functionalize the surface, first cleaved the mica surface by etching, then we incubated it for 1 minute with 10 µl of 0.1% (v/v) 3-aminopropyl-triethoxysilane (APTES; Sigma Aldrich) in Milli-Q water. The substrate was rinsed three times with 1 mL of Milli-Q water and dried by a gentle stream of nitrogen gas. Subsequently, 10 µl 0.3 ng/µl of DNA and NX11-DNA complexes in 1.5 mM Tris-HCl pH 8.0 was deposited onto the positive functionalized surface. The droplet was incubated for 10 min, rinsed by 1 mL of Milli-Q water and dried by a gentle stream of nitrogen gas. The entire preparation process was conducted at room temperature and under laminar flow.

AFM maps of 3D morphology were acquired for both samples in a regime of constant phase change, with 2 nm/pixel resolution by means of a Multimode-8 (Bruker) operating in tapping mode and equipped with a gold coated probe (HQ:NSC14/Cr-Au BS, 5 N/m; Bruker) with a nominal radius < 8 nm.

Scanning probe image processor (version 6.7.3, Image Metrology, Denmark) software was used for image processing and further analysis. Cross sectional profiles of both DNA and NX11-DNA were extracted for statistical analysis at the level of individual molecules (n=35). Additionally, a FFT of the cross-sectional profile and the gradient of the cross-sectional profile were used to derive variations in periodicity for NX11-DNA molecules (n=10), allowing for observation of higher and smaller periodicity components.

### Negative stain electron microscopy collection and processing

NX11-DNA were diluted to 12 ng/μL DNA concentration prior to application of 3 μL of sample onto freshly glow-discharged 400-mesh copper grids (Ted Pella). Sample was incubated on the grid for 15 - 30 sec before excess liquid blotted away with filter paper (Whatman). 3 μL of 2% w/v uranyl formate (UF) stain was applied to the grid and immediately blotted away before an additional 3 μL of UF stain was applied. Stain was blotted off by filter paper, and a final 3 μL of UF stain was applied and allowed to incubate for ∼30 sec. Finally, the stain was blotted away and the grids were allowed to dry for 3 min. Prepared grids were imaged using EPU 2.0 on a 120 kV Talos L120C transmission electron microscope (Thermo Fisher Scientific) at 72,000× magnification with a BM-Ceta camera. Data processing was done in CryoSPARC^47^, starting with CTF correction, particle picking, and 2D classification.

### Cryo-electron microscopy data collection, processing, and analysis

CryoEM grids were prepared by applying 3.5 μL of 0.6 mg/mL NX11Q21.5-DNA48 in TBS with 2.5% glycerol to glow-discharged 400 mesh C-flat grids (CF-2/2-4C, 2 μm holes, 2 μm spacing; Electron Microscopy Sciences). The grids were blotted with a blot force of 0 for 6.5 seconds at 100% humidity and 20 °C, followed by plunge-freezing in liquid ethane using a Vitrobot Mark IV (FEI Thermo Scientific). Grids were screened and collected using a Titan Krios transmission electron microscope (FEI Thermo Scientific) operated at 300 kV, equipped with a K3 Summit direct detector and energy filter, located at the Janelia HHMI research campus.

Automated data collection, totaling 10,595 movies, was performed using SerialEM software with a pixel size of 0.4135 Å in super-resolution mode. Each movie was fractionated into 75 frames, with a total dose of ∼60 e-/Å^2^. Data were processed in CryoSPARC, beginning with patch motion correction and CTF estimation using default parameters. Approximately 1.2 million particles were picked using the filament tracer, which were subsequently sorted into 150 2D class averages. Both type I and type II fiber classes were identified, and their power spectra were averaged to measure the helical pitch of each fiber class. Despite multiple attempts at 3D reconstructions, the resulting models lacked the necessary resolution and confidence for model building or detailed interpretation and were thus excluded from our analysis.

### Antigen-adjuvant formulations and immunizations in mice

NX11-immunogens (NX11-majorX3, NX11-majorX5, NX11-minorX3, NX11-minorX5, NX11-juncX3, NX11-juncX5) were assembled with pharmaceutical grade DNA_250_ (NoLimits, Thermo Fisher Scientific) at a DNA concentration of 10.33 ng/μL and 1.1-fold excess of protein (assuming 1 NX protein binding to 1 bp) in TBS + 5% glycerol. Complexes were incubated overnight at room temperature, split in 12 times 5 µL and flash frozen in -80 °C.

Directly before injection I53_dn5_majorX5 and NX11-DNA immunogen aliquots were thawed on ice and 1.5 µg was mixed with Addavax adjuvant in a 1:1 ratio. This vaccine formulation (in a volume of 100 uL) was injected intramuscularly in the hind legs of the mice (5 mice per group).

### I53_dn5_majorX5-L1 Protein expression and purification

I53_dn5A.1_majorx5-L1 component containing a His-tag was expressed in a pET29b+ vector in BL21(DE3) E. coli. Inoculated cultures were expressed in Terrific Broth II (MP Biomedicals) in 2L baffled flasks at 37°C and 220 RPM until OD600 reached 0.6-0.8. Cultures were then induced with 1 mM IPTG, and temperature was lowered to 18°C and grown for 16-18h. Cells were harvested by centrifugation and lysed by high-pressure microfluidization.

Clarified *E. coli* lysate was then applied to Nickel-NTA gravity (QIAGEN) flow column that was previously equilibrated with 50 mM Tris, 150 mM NaCl, 20 mM Imidazole pH 8.0 (Buffer A). After sample application, the column was washed with 5 CV of Buffer A, and then with 10 CV of Buffer A + 0.75% CHAPS for endotoxin removal, and once again with 5 CV of Buffer A. Protein was eluted with 50 mM Tris, 150 mM NaCl, 300 mM Imidazole pH 8.0 (Buffer B). The eluted fractions were then analyzed by SDS-PAGE to confirm the presence and purity of I53_dn5A.1_majorx5-L1. IMAC purified protein was then further purified by SEC using a Superdex 200 Increase 10/300 (Cytiva) equilibrated in 50 mM Tris, 150 mM NaCl, 5% Glycerol. Fractions were collected, pooled, and analyzed by SDS-PAGE to confirm purity, UV/vis spectroscopy to obtain protein concentration, and Limulus amebocyte lysate assay (Charles River) for endotoxin.

### Enzyme-linked immunosorbent assay (ELISA) for measuring binding antibody titers

96-well flat-bottom Immuno MaxiSorp plates were coated with 20 ng of *pf*CSP antigen per well in 100 µL of PBS buffer overnight at 4°C. Coated plates were washed 3x with PBS with 0.05% Tween-20 (ELISA wash buffer). Plates were blocked with 5% Casein in ELISA wash buffer (blocking buffer) for 1 hour at room temperature and plates were subsequently washed 3x with ELISA wash buffer. Mouse sera was individually diluted 1:100 in blocking buffer and was then diluted 4-fold down the plate to a final dilution of 1.05E+08 and incubated at room temperature for 45 minutes. Plates were washed and then Goat anti-mouse IgG-HRP was diluted to 1:2000 in ELISA wash buffer and added to the plates for 45 minutes at room temperature. Plates were washed a final time and 100 µL of TMB 1-component peroxidase substrate (SeraCare) was added for 5 minutes and subsequently neutralized with 2N HCl. Absorbance at 450 nm was measured with a Neo2 plate reader to determine endpoint values.

### *In vivo* immunogenicity

Female B6(Cg)-*Tyrc-2J*/J mice were purchased from The Jackson Laboratory (strain code 000058) at 6 weeks of age. Mice were housed in a specific-pathogen free facility within the Department of Comparative Medicine at the University of Washington, Seattle, accredited by the Association for Assessment and Accreditation of Laboratory Animal Care (AAALAC). Animal studies were conducted in accordance with the University of Washington’s Institutional Animal Care and Use Committee under protocol 4470-01. For each immunization, low-endotoxin immunogens were diluted in buffer and mixed with 1:1 v/v AddaVax adjuvant (InvivoGen vac-adx-10) to obtain a final dose of 1.5 μg of immunogen per animal, per injection. At 8 weeks of age, 5 mice per group were injected intramuscularly in the quadriceps with 50 μL of immunogen per hind leg at weeks 0 and 3. Animals were bled using the submental route at weeks 2 & 5. Whole blood was collected in serum separator tubes (BD #365967) and rested for 30 min at room temperature for coagulation. Tubes were then centrifuged for 10 min at 2,000 x *g* and serum was collected and stored at -80°C until use.

## Acknowledgements

This work was supported financially by the VLAG Graduate School Research Fellowship and a Fulbright Visiting Scholar Fellowship to R.J.dH, the Bill & Melinda Gates Foundation (INV-010680), and The Audacious Project at the Institute for Protein Design. The authors thank Geoff Hutchinson for assistance with ELISA assays, and Drs. Marie Pancera and Nick Hurlburt for providing PfCSP antigen. We thank the HHMI Janelia CryoEM Facility staff for help in microscope operation and data collection.

