## Supplementary Information for "Computational redesign of TALE proteins for DNA-templated assembly of protein fibers"

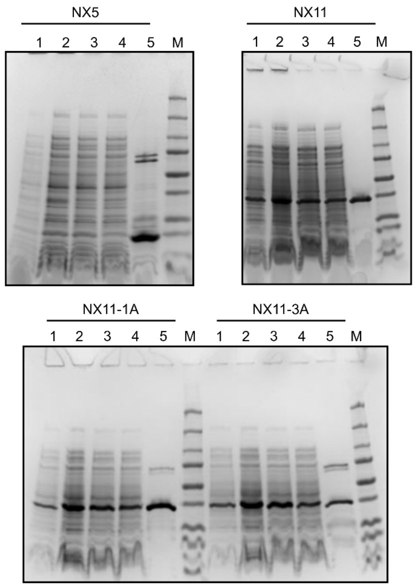


**Figure S1. SDS-PAGE of purification of NX5, NX11, NX11-1A, and NX11-3A proteins from *E. coli.*** 1) Induced cell pellet, 2) cell lysate, 3) clarified lysate, 4) IMAC flow-through, and 5) IMAC eluate. M) Precision Plus Protein Dual Color Standard (250 kDa, 150 kDa, 100 kDa, 75 kDa, 50 kDa, 37 kDa, 20 kDa, 15 kDa, 10 kDa; Bio-Rad).

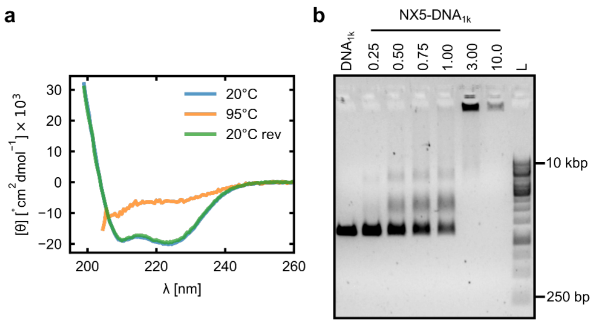


**Figure S2. Biochemical characterization and assembly of NX5 with DNA.**

**(a)** Circular Dichroism of purified NX5 in PBS. Molar residue ellipticity [θ] is plotted as a function of wavelength. NX5 was largely unfolded at 95 °C, but upon cooling back to 20 °C (20 °C rev) re-folded to its original conformation. **(b)** EMSA showing co-assembly of NX5 with 15 ng/µL $DNA_{1k}$ at various NX5:$DNA_{1k}$ ratios. Due to charge neutralization and increased size, the NX5-DNA complexes migrated slower, indicating assembly. The $DNA_{1k}$ lane contained a DNA-only sample. L was a 1 kb gene ruler (ThermoFisher) with minimal and maximal size indicated.

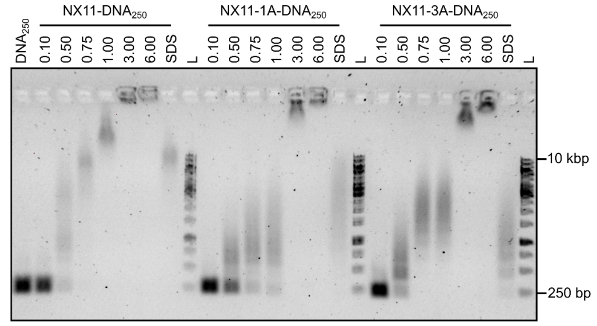


**Figure S3. EMSA comparing co-assembly of NX11, NX11-1A, and NX11-3A with $DNA_{250}$ at various ratios of protein to DNA.**

All three proteins fully assembled at excess above 3.00. However, alanine interface mutants NX11-1A and NX11-3A appeared to shift at higher excess of protein:DNA, indicating that weaker interfaces have less cooperative assembly. The “SDS” lane contained a 3-fold excess assembly of NX11-DNA loaded in presence of ~0.2% SDS to denature the complexes. In all cases the complexes were not fully denatured as they appear to smear on the gel. “L” was a 1 kb gene ruler (ThermoFisher) with minimal and maximal size indicated.

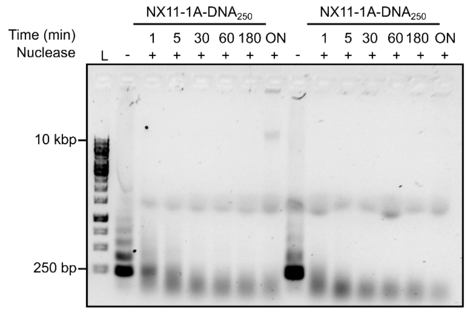


**Figure S4. Benzonase assay of NX11-1A and NX11-3A assembled with $DNA_{250}$.**
Benzonase nuclease was added to the protein-DNA complexes in reaction buffer (10 mM Tris pH 7.5, 2.5 mM $MgCl_2$, 0.1 mM $CaCl_2$) and incubated for 1 min, 5 min, 30 min, 60 min, 180 min, and overnight (ON). A control DNA-only sample lacking Benzonase was loaded as reference. Benzonase cleavage was stopped by addition of a stopping buffer (50 mM EDTA pH 7.4). Samples were loaded in the presence of 0.2% SDS to fully denature the complexes and observe if the 250bp template DNA was cleaved. This appeared to be the case for both NX11-1A and NX11-3A: compared to the DNA-only sample, less intense bands were observed and the DNA was shorter than 250bp. “L” was a 1 kb gene ruler (ThermoFisher) with minimal and maximal size indicated.

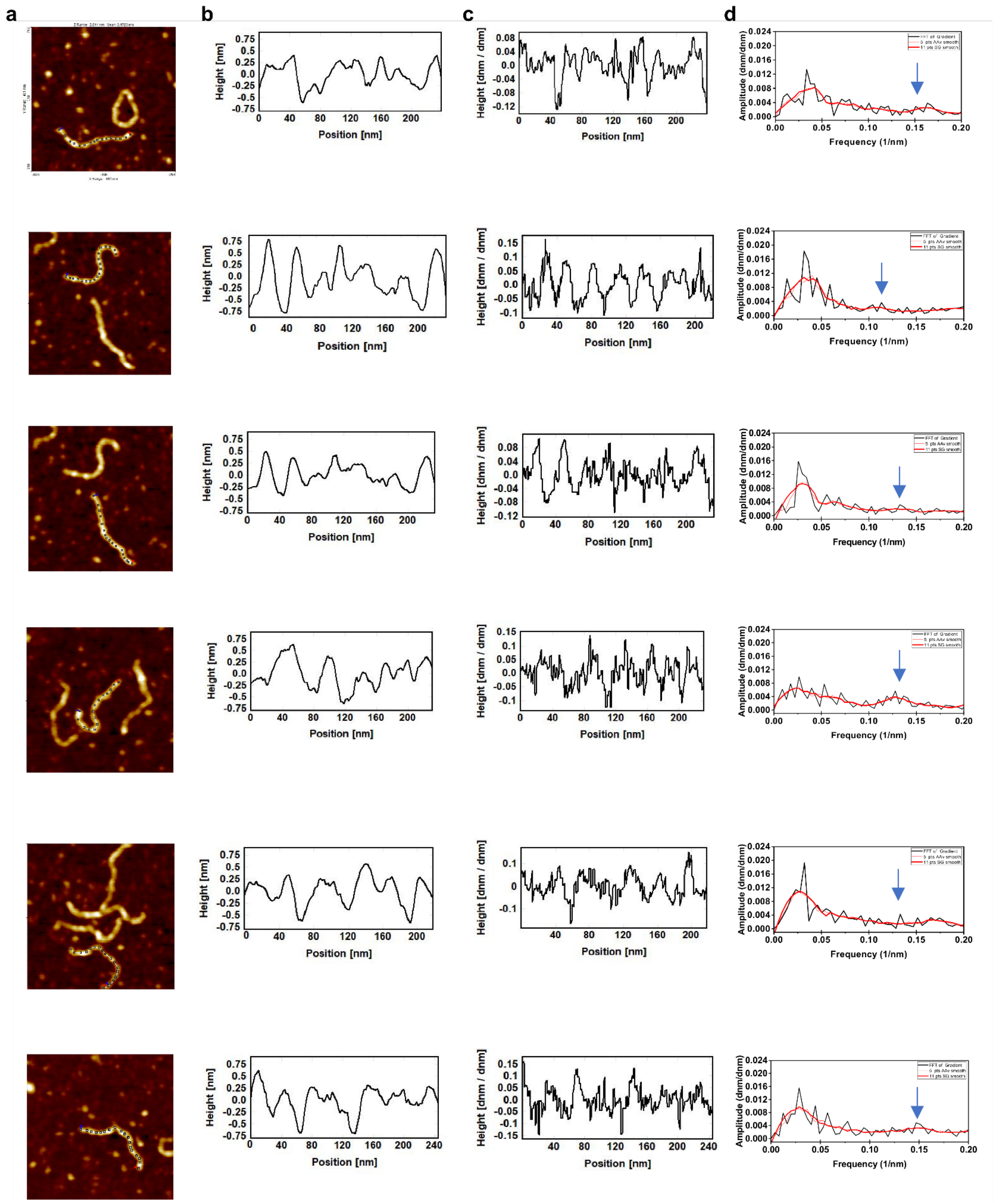


**Figure S5. AFM periodicity analysis traces of 6 representative NX11-DNA particles.**
**(a)** AFM morphology maps, **(b)** line profiles, **(c)** gradients of the line profiles, and **(d)** fourier transforms of the gradient profiles. Blue arrow indicates the minor periodicity corresponding to ~7 nm.

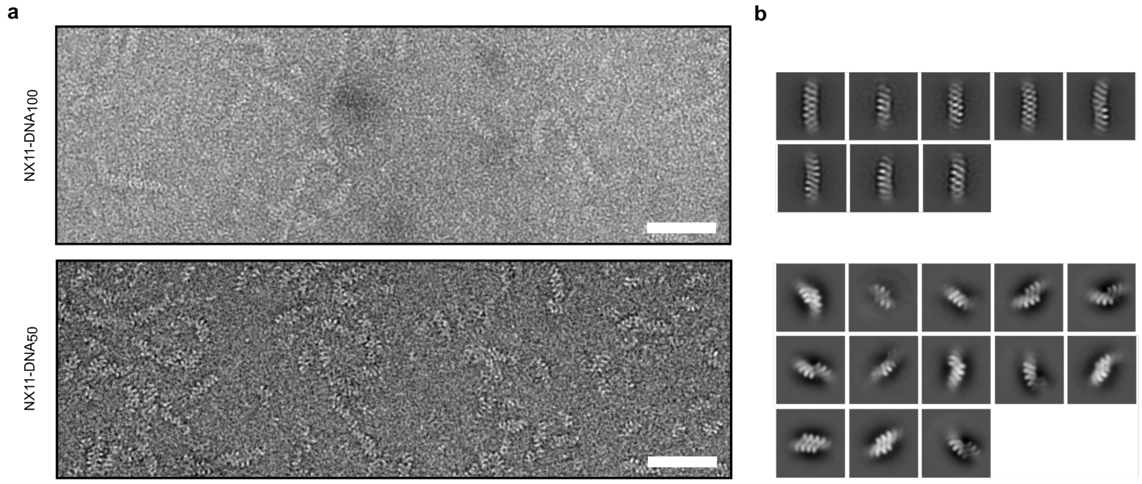


**Figure S6. nsEM of NX11-DNA$_{100}$ and NX11-DNA$_{50}$ complexes.**
**(a)** Representative negatively stained electron micrographs. Both assemblies produce fibrous complexes of the approximate expected contour length. **(b)** 2D class averages demonstrate that in the case of shorter DNA lengths (< 100 bp), NX11-DNA complexes tend to stack side-by-side as can be seen from a selected subset of 2D class averages. In the case of DNA > 100 bp, complexes do not stack side-by-side, likely because they are less rigid. Scale bars: 50 nm.

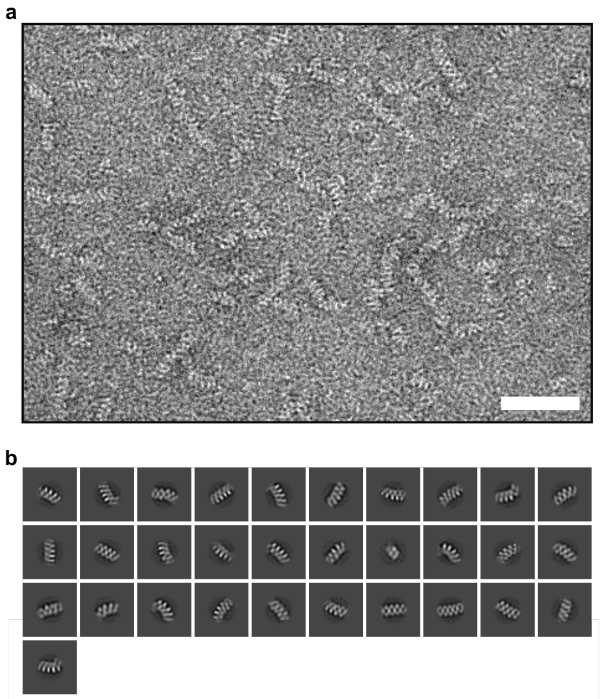


**Figure S7. nsEM of NX11$^{21Q.5}$-DNA$_{50}$ complexes.**
**(a)** Negatively stained electron micrographs. Scale bar: 50 nm. **(b)** Selected 2D class averages.

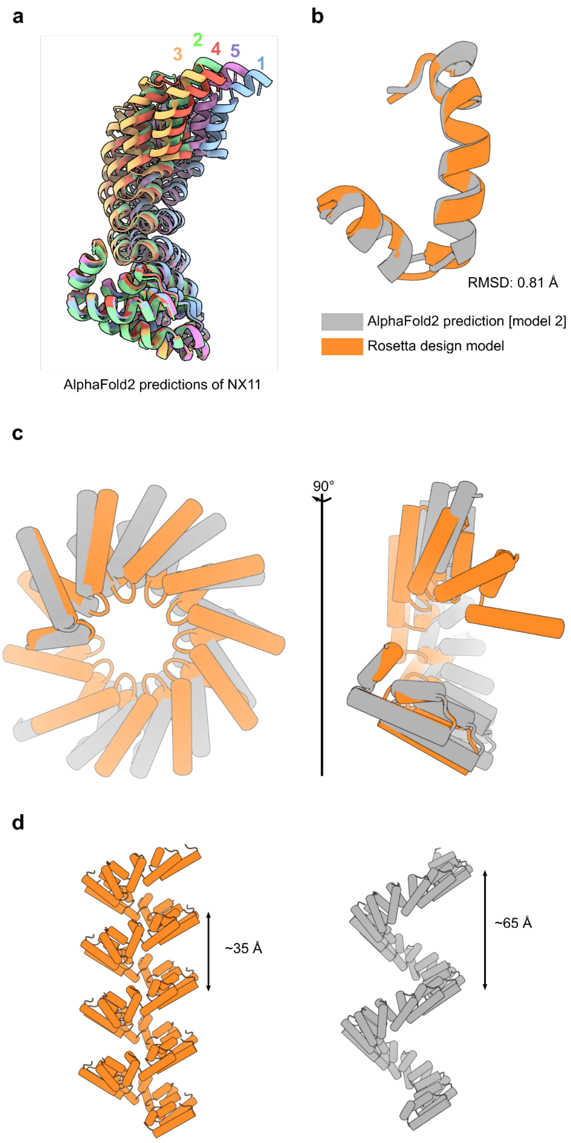


**Fig S8. Comparison of NX11 Rosetta design model and AlphaFold2 predictions.**
**(a)** AlphaFold2 predictions of NX11 with all 5 parameter models. The different AF2 models show some plasticity with respect to the apparent pitch of the NX11. **(b)** Alignment of a single repeat from from NX11, demonstrating that at the single repeat level the AF2 model 2 prediction agrees with the Rosetta model. **(c)** Alignment of the first repeat of the NX11 protein. **(d)** Repeat propagation using helical symmetry, where the symmetry parameters are inferred from two neighboring repeats in the design model or the AF2 prediction. Here the difference in helical rise in the predicted AF2 model 2 becomes apparent. The Rosetta design model follows the DNA helix rise of ~34 Å, but the AF2 model 2 predicts an extended rise of ~65 Å.

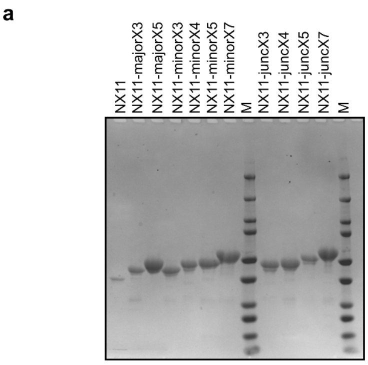


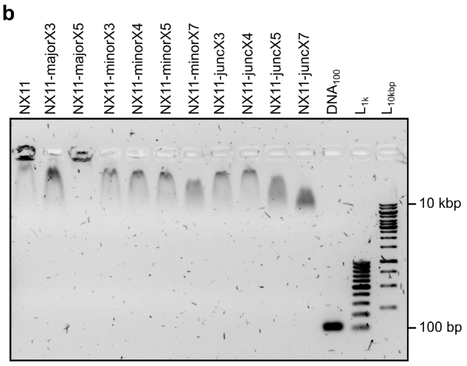


**Figure S9. Biochemical characterization of epitope-bearing NX11 variants.**
**(a)** SDS-PAGE of NX11 variants displaying various malaria antigens (major, minor, or junctional) at different repeat numbers. NX11 carrying no antigens was loaded as a reference. **(b)** EMSA of NX11 variants displaying malaria antigens assembled with $DNA_{100}$ (NoLimits, Thermo Fisher Scientific) at 3-fold excess. In all cases the DNA migration was slower due to assembly of protein. "$L_{1k}$" is a 100 bp gene ruler (ThermoFisher) and "$L_{10kbp}$" is a 1 kbp gene ruler (ThermoFisher) with maximal and minimal sizes indicated.

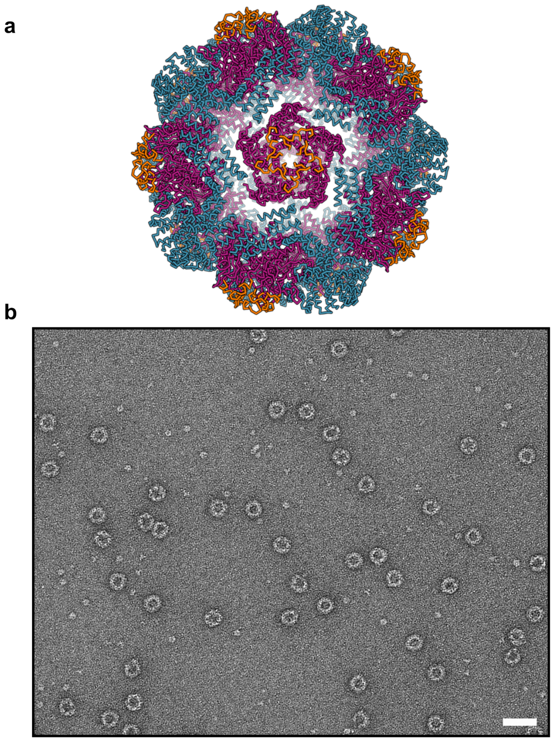

**Figure S10. Design model and electron micrographs of I53_dn5_majorX5.**
**(a)** Design model of I53_dn5_majorX5 with I53_dn5B highlighted in blue, I53_dn5A highlighted in purple, and majorX5 insertion highlighted in orange. Each particle displays 60 copies of the majorX5 antigen. **(b)** Representative negative stain electron micrograph of *in vitro* assembled I53_dn5_majorX5. Scale bar: 50 nm.

**Table S1. Novel protein sequences generated in this study.**

| Design | Protein sequence |
|---|---|
| NX11 | MGHHHHHHGSSENLYFQGSMPDQVVAIAKRGGKQALETVQRLLPELTQKHGLTPDQVVAIAKRGGKQALETVQRLLPELTQKHGLTPDQVVAIAKRGGKQALETVQRLLPELTQKHGLTPDQVVAIAKRGGKQALETVQRLLPELTQKHGLTPDQVVAIAKRGGKQALETVQRLLPELTQKHGLTPDQVVAIAKRGGKQALETVQRLLPELTQKHGLTPDQVVAIAKRGGKQALETVQRLLPELTQKHGLTPDQVVAIAKRGGKQALETVQRLLPELTQKHGLTPDQVVAIAKRGGKQALETVQRLLPELTQKHGLTPDQVVAIAKRGGKQALETVQRLLPELTQKHGLTPDQVVAIAKRGGKQALETVQRLLPELTQKHGW |
| NX5 | MGHHHHHHGSSENLYFQGSMPDQVVAIAKRGGKQALETVQRLLPELTQKHGLTPDQVVAIAKRGGKQALETVQRLLPELTQKHGLTPDQVVAIAKRGGKQALETVQRLLPELTQKHGLTPDQVVAIAKRGGKQALETVQRLLPELTQKHGLTPDQVVAIAKRGGKQALETVQRLLPELTQKHGW |
| NX11-1A | MGHHHHHHGSSENLYFQGSMPDQVAAIAKRGGKQALETVQRLLPELTQKHGLTPDQVVAIAKRGGKQALETVQRLLPELTQKHGLTPDQVVAIAKRGGKQALETVQRLLPELTQKHGLTPDQVVAIAKRGGKQALETVQRLLPELTQKHGLTPDQVVAIAKRGGKQALETVQRLLPELTQKHGLTPDQVVAIAKRGGKQALETVQRLLPELTQKHGLTPDQVVAIAKRGGKQALETVQRLLPELTQKHGLTPDQVVAIAKRGGKQALETVQRLLPELTQKHGLTPDQVVAIAKRGGKQALETVQRLLPELTQKHGLTPDQVVAIAKRGGKQALETVQRLLPELTQKHGLTPDQVVAIAKRGGKQALETVQRLLPELTQKHGW |
| NX11-3A | MGHHHHHHGSSENLYFQGSMPDQVAAIAKRGGKQAAETVQRLLPELTQKHGLTPDQVVAIAKRGGKQALETVQRLLPELTQKHGLTPDQVVAIAKRGGKQALETVQRLLPELTQKHGLTPDQVVAIAKRGGKQALETVQRLLPELTQKHGLTPDQVVAIAKRGGKQALETVQRLLPELTQKHGLTPDQVVAIAKRGGKQALETVQRLLPELTQKHGLTPDQVVAIAKRGGKQALETVQRLLPELTQKHGLTPDQVVAIAKRGGKQALETVQRLLPELTQKHGLTPDQVVAIAKRGGKQALETVQRLLPELTQKHGLTPDQVVAIAKRGGKQALETVQRLLPELTQKHGLTPDQVVAIAKRGGKQALETVQRALPELTQKHGW |
| NX11-21.Q5 | MGHHHHHHGSSENLYFQGSMPDQVVAIAKRGGKQALETVQRLLPELTQKHGLTPDQVVAIAKRGGKQALETVQQLLPELTQKHGLTPDQVVAIAKRGGKQALETVQRLLPELTQKHGLTPDQVVAIAKRGGKQALETVQQLLPELTQKHGLTPDQVVAIAKRGGKQALETVQRLLPELTQKHGLTPDQVVAIAKRGGKQALETVQQLLPELTQKHGLTPDQVVAIAKRGGKQALETVQRLLPELTQKHGLTPDQVVAIAKRGGKQALETVQQLLPELTQKHGLTPDQVVAIAKRGGKQALETVQRLLPELTQKHGLTPDQVVAIAKRGGKQALETVQQLLPELTQKHGLTPDQVVAIAKRGGKQALETVQRLLPELTQKHGW |
| NX11-majorX3 | MGHHHHHHGSSENLYFQGSMPDQVVAIAKRGGKQALETVQRLLPELTQKHGLTPDQVVAIAKRGGKQALETVQRLLPELTQGNANPNANPNANPGDQVVAIAKRGGKQALETVQRLLPELTQKHGLTPDQVVAIAKRGGKQALETVQRLLPELTQKHGLTPDQVVAIAKRGGKQALETVQRLLPELTQGNANPNANPNANPGDQVVAIAKRGGKQALETVQRLLPELTQKHGLTPDQVVAIAKRGGKQALETVQRLLPELTQKHGLTPDQVVAIAKRGGKQALETVQRLLPELTQKHGLTPDQVVAIAKRGGKQALETVQRLLPELTQGNANPNANPNANPGDQVVAIAKRGGKQALETVQRLLPELTQKHGW |
| NX11-majorX5 | MGHHHHHHGSSENLYFQGSMPDQVVAIAKRGGKQALETVQRLLPELTQKHGLTPDQVVAIAKRGGKQALETVQRLLPELTQGNANPNANPNANPNANPNANPGDQVVAIAKRGGKQALETVQRLLPELTQKHGLTPDQVVAIAKRGGKQALETVQRLLPELTQKHGLTPDQVVAIAKRGGKQALETVQRLLPELTQGNANPNANPNANPNANPNANPGDQVVAIAKRGGKQALETVQRLLPELTQKHGLTPDQVVAIAKRGGKQALETVQRLLPELTQ |

| | |
|---|---|
| | KHGLTPDQVVAIAKRGGKQALETVQRLLPELTQKHGLTPDQVVAIAKRGGKQALETVQRLLPELTQGNANPNANPNANPNANPNANPGDQVVAIAKRGGKQALETVQRLLPELTQKHGW |
| NX11-minorX3 | MGHHHHHHGSSENLYFQGSMPDQVVAIAKRGGKQALETVQRLLPELTQKHGLTPDQVVAIAKRGGKQALETVQRLLPELTQKHGLTPDQVVAIAKRGGKQALETVQRLLPELTQGNPNVDPNANPNVGDQVVAIAKRGGKQALETVQRLLPELTQKHGLTPDQVVAIAKRGGKQALETVQRLLPELTQKHGLTPDQVVAIAKRGGKQALETVQRLLPELTQGNPNVDPNANPNVGDQVVAIAKRGGKQALETVQRLLPELTQKHGLTPDQVVAIAKRGGKQALETVQRLLPELTQKHGLTPDQVVAIAKRGGKQALETVQRLLPELTQKHGLTPDQVVAIAKRGGKQALETVQRLLPELTQGNPNVDPNANPNVGDQVVAIAKRGGKQALETVQRLLPELTQKHGW |
| NX11-minorX4 | MGHHHHHHGSSENLYFQGSMPDQVVAIAKRGGKQALETVQRLLPELTQKHGLTPDQVVAIAKRGGKQALETVQRLLPELTQKHGLTPDQVVAIAKRGGKQALETVQRLLPELTQGNPNVDPNANPNVDPNAGDQVVAIAKRGGKQALETVQRLLPELTQKHGLTPDQVVAIAKRGGKQALETVQRLLPELTQKHGLTPDQVVAIAKRGGKQALETVQRLLPELTQGNPNVDPNANPNVDPNAGDQVVAIAKRGGKQALETVQRLLPELTQKHGLTPDQVVAIAKRGGKQALETVQRLLPELTQKHGLTPDQVVAIAKRGGKQALETVQRLLPELTQKHGLTPDQVVAIAKRGGKQALETVQRLLPELTQGNPNVDPNANPNVDPNAGDQVVAIAKRGGKQALETVQRLLPELTQKHGW |
| NX11-minorX5 | MGHHHHHHGSSENLYFQGSMPDQVVAIAKRGGKQALETVQRLLPELTQKHGLTPDQVVAIAKRGGKQALETVQRLLPELTQKHGLTPDQVVAIAKRGGKQALETVQRLLPELTQGNPNVDPNANPNVDPNANPNVGDQVVAIAKRGGKQALETVQRLLPELTQKHGLTPDQVVAIAKRGGKQALETVQRLLPELTQKHGLTPDQVVAIAKRGGKQALETVQRLLPELTQGNPNVDPNANPNVDPNANPNVGDQVVAIAKRGGKQALETVQRLLPELTQKHGLTPDQVVAIAKRGGKQALETVQRLLPELTQKHGLTPDQVVAIAKRGGKQALETVQRLLPELTQKHGLTPDQVVAIAKRGGKQALETVQRLLPELTQGNPNVDPNANPNVDPNANPNVGDQVVAIAKRGGKQALETVQRLLPELTQKHGW |
| NX11-minorX7 | MGHHHHHHGSSENLYFQGSMPDQVVAIAKRGGKQALETVQRLLPELTQKHGLTPDQVVAIAKRGGKQALETVQRLLPELTQKHGLTPDQVVAIAKRGGKQALETVQRLLPELTQGNPNVDPNANPNVDPNANPNVDPNANPNVGDQVVAIAKRGGKQALETVQRLLPELTQKHGLTPDQVVAIAKRGGKQALETVQRLLPELTQKHGLTPDQVVAIAKRGGKQALETVQRLLPELTQGNPNVDPNANPNVDPNANPNVDPNANPNVGDQVVAIAKRGGKQALETVQRLLPELTQKHGLTPDQVVAIAKRGGKQALETVQRLLPELTQKHGLTPDQVVAIAKRGGKQALETVQRLLPELTQKHGLTPDQVVAIAKRGGKQALETVQRLLPELTQGNPNVDPNANPNVDPNANPNVDPNANPNVGDQVVAIAKRGGKQALETVQRLLPELTQKHGW |
| NX11-juncX3 | MGHHHHHHGSSENLYFQGSMPDQVVAIAKRGGKQALETVQRLLPELTQKHGLTPDQVVAIAKRGGKQALETVQRLLPELTQKHGLTPDQVVAIAKRGGKQALETVQRLLPELTQGNPDPNANPNVDPNAGDQVVAIAKRGGKQALETVQRLLPELTQKHGLTPDQVVAIAKRGGKQALETVQRLLPELTQKHGLTPDQVVAIAKRGGKQALETVQRLLPELTQGNPDPNANPNVDPNAGDQVVAIAKRGGKQALETVQRLLPELTQKHGLTPDQVVAIAKRGGKQALETVQRLLPELTQKHGLTPDQVVAIAKRGGKQALETVQRLLPELTQKHGLTPDQVVAIAKRGGKQALETVQRLLPELTQGNPDPNANPNVDPNAGDQVVAIAKRGGKQALETVQRLLPELTQKHGW |
| NX11-juncX4 | MGHHHHHHGSSENLYFQGSMPDQVVAIAKRGGKQALETVQRLLPELTQKHGLTPDQVVAIAKRGGKQALETVQRLLPELTQKHGLTPDQVVAIAKRGGKQALETVQRLLPELTQGNPDPNANPNVDPNANPNVGDQVVAIAKRGGKQALETVQRLLPELTQKHGLTPDQVVAIAKRGGKQALETVQRLLPELTQKHGLTPDQVVAIAKRGGKQALETVQRLLPELTQGNPDPNANPNVDPNANPNVGDQVVAIAKRGGKQALETVQRLLPELTQKHGLTPDQVVAIAKRGGKQALETVQRLLPELTQKHGLTPDQVVAIAKRGGKQALETVQRLLPELTQKHGLTPDQVVAIAKRGGKQALETV |

| | |
|---|---|
| | QRLLPELTQGNPDPNANPNVDPNANPNVGDQVVAIAKRGGKQALETVQRLLPELTQKHGW |
| NX11-juncX5 | MGHHHHHHGSSENLYFQGSMPDQVVAIAKRGGKQALETVQRLLPELTQKHGLTPDQVVAIAKRGGKQALETVQRLLPELTQKHGLTPDQVVAIAKRGGKQALETVQRLLPELTQGNPDPNANPNVDPNANPNVDPNAGDQVVAIAKRGGKQALETVQRLLPELTQKHGLTPDQVVAIAKRGGKQALETVQRLLPELTQKHGLTPDQVVAIAKRGGKQALETVQRLLPELTQGNPDPNANPNVDPNANPNVDPNAGDQVVAIAKRGGKQALETVQRLLPELTQKHGLTPDQVVAIAKRGGKQALETVQRLLPELTQKHGLTPDQVVAIAKRGGKQALETVQRLLPELTQKHGLTPDQVVAIAKRGGKQALETVQRLLPELTQGNPDPNANPNVDPNANPNVDPNAGDQVVAIAKRGGKQALETVQRLLPELTQKHGW |
| NX11-juncX7 | MGHHHHHHGSSENLYFQGSMPDQVVAIAKRGGKQALETVQRLLPELTQKHGLTPDQVVAIAKRGGKQALETVQRLLPELTQKHGLTPDQVVAIAKRGGKQALETVQRLLPELTQGNPDPNANPNVDPNANPNVDPNANPNVDPNAGDQVVAIAKRGGKQALETVQRLLPELTQKHGLTPDQVVAIAKRGGKQALETVQRLLPELTQKHGLTPDQVVAIAKRGGKQALETVQRLLPELTQGNPDPNANPNVDPNANPNVDPNANPNVDPNAGDQVVAIAKRGGKQALETVQRLLPELTQKHGLTPDQVVAIAKRGGKQALETVQRLLPELTQKHGLTPDQVVAIAKRGGKQALETVQRLLPELTQKHGLTPDQVVAIAKRGGKQALETVQRLLPELTQGNPDPNANPNVDPNANPNVDPNANPNVDPNAGDQVVAIAKRGGKQALETVQRLLPELTQKHGW |

**Table S2. PfCSP epitopes used in the mice study.**

| Immunogen | Epitope sequence |
|---|---|
| majorX3 | NANPNANPNANP |
| majorX5 | NANPNANPNANPNANPNANP |
| minorX3 | NPNVDPNANPNV |
| minorX5 | NPNVDPNANPNVDPNANPNV |
| juncX3 | NPDPNANPNVDPNA |
| juncX5 | NPDPNANPNVDPNANPNVDPNA |
